# Induction of the HIF pathway: Differential regulation by chemical hypoxia and oxygen glucose deprivation

**DOI:** 10.1101/525006

**Authors:** Alicia E. Novak, Susan M. Jones, J. Paul Elliott

## Abstract

The Hypoxia Inducible Factor (HIF) proteins are the master regulators in the cellular response to varying oxygen levels, including hypoxia. The HIF complex is stabilized and accumulates when oxygen levels drop through inhibition of a degradative enzyme. An active HIF complex can act as a transcriptional regulator of hundreds of genes. In turn, these genes determine the response of the cell by inducing pathways which can promote survival, or result in cell death. However, little is known about the regulation of the transcriptional process. We were interested in learning more about the time dependence of transcriptional activation in order to target those pathways which could enhance cell survival after ischemia. Using mouse hippocampal organotypic cultures (HOTCs), we compared oxygen-glucose deprivation with the hypoxia mimetic cobalt, which inhibits the oxygen dependent prolyl hydroylase and blocks degradation of the HIF proteins. We demonstrated that two of the most studied HIF target genes (VEGF, EPO) as well as HIF structural genes show complex time and dose-dependent expression patterns in response to the two different insults. Understanding of these molecular responses is crucial for the development of future treatments to enhance recovery from hypoxia and stroke.

## Introduction

An adaptive response to a decrease in oxygen levels in humans occurs when an individual has reduced blood flow in a region of their brain, as in stroke and traumatic brain injury. The resulting focal brain ischemia initiates a complex altered state, in which there are rapid molecular changes in gene and protein composition that influence the viability of the damaged tissue and cells immediately and many hours after the initial insult. This process is not well understood and has been the focus of much research to advance stroke and TBI treatments.

HIF (hypoxia inducible factor) is a well-studied protein complex that forms when oxygen levels decrease [1, 2]. Pathways involved in angiogenesis, glucose metabolism, erythropoiesis and others contain genes that are regulated by the HIF pathway [3–5]. During normoxia, the HIF alpha subunit protein (HIFα) is hydroxylated by a prolyl hydroxylase enzyme (PHD) which targets it for degradation by the von Hippel Lindau E3 ubiquitin ligase (VHL) [6, 7]. A subsequent drop in oxygen inactivates the PHD molecule allowing stabilization of HIFα and further dimerization with a HIFβ subunit to form a functional protein complex [6–8]. The stable HIFα/β dimer, in conjunction with other co-factors such as p300/Creb binding protein, then binds to a hypoxic response element (HRE) on target genes to cause rapid transactivation [9–11]. Compounds which interfere with the targeted degradation of HIFα subunits can mimic hypoxia. Cobalt Chloride (CoCl_2_) is a heavy metal which has been reported to interfere with hydroxylation of the Hifα subunit by PHD and subsequent binding by VHL [12, 13].

At the gene level, three alpha subunits (Hif1α, Epas1or Hif2α and Hif3α) and two beta subunits (Hif1β/ARNT and Hif2β/ARNT2), each encoding a functional protein, have been identified in mammals [14–18]. While HIF1α is the most abundant and well described alpha subunit, both Hif2α and Hif3α are also expressed in the brain in a cell and region specific manner [14, 17, 19–22]. Transcriptional regulation of both Hif1α and Hif3α, but not Hif2α, has been reported with hypoxic induction [19, 23–26]. HIFβ subunits are more ubiquitously expressed in the brain and are more resistant to hypoxic regulation at the transcript level [19, 27, 28]. Establishing the transcriptional regulation of these genes in a single model is important for understanding the activation of compensatory mechanisms and molecular alterations in the brain after an oxygen challenge. While individual bodies of research have looked at the hypoxic induction of each of the HIF structural subunits, a systematic approach of looking at the transcriptional expression of all HIF subunit genes and target genes in a single model during a time course after a hypoxic event with or without glucose has not been established.

In this work, we used a mouse hippocampal organotypic culture (HOTC) to elucidate transcriptional changes in all HIF structural subunits by induction of the HIF pathway. To assess the activation of the HIF pathway in our model, we first characterized the expression of two well-known HIF targets, vascular endothelial growth factor (Vegf) and erythropoietin (Epo). Three Vegf genes exist, however, only Vegfa (also known as Vegf) has been shown to be regulated by hypoxia [29]. HOTCs are a good model to study the ischemic and hypoxic pathways as they closely mimic conditions found *in vivo* [30]. HIF induction was studied using both OGD, a well described method for HOTCs, and CoCl_2_, a hypoxic mimetic whose use in HOTCs has not been described in the literature. Each model was chosen to provide comparative information about activation of the HIF pathway; OGD by an acute and transient oxygen decrease accompanied by a loss of glucose (OGD), and CoCl_2_ to pharmacologically mimic chronic hypoxia with glucose.

We demonstrated that there is a complex and specific regulation of Vegf and Epo genes and HIF structural subunit genes that was duration and dose dependent. While Vegf and Hif3α changes were induced by both treatments, differential increases in transcription of Epo (only with CoCl_2_) and Hif1α (only with OGD) were seen. These results provide new information about the transcriptional regulation of HIF target genes and structural subunits after a drop of oxygen levels in the HOTC system.

## Materials and Methods

All studies were conducted using NIH guidelines for animal care and use. All protocols were approved by the Swedish Medical Center Institutional Animal Care and Use Committee (IACUC).

### Hippocampal cultures

Mouse organotypic hippocampal cultures (HOTC) were made as previously described [31]. Briefly, 7 day old C57BL/6NHsd mice (Harlan Laboratories) were sacrificed using decapitation as per American Veterinarian Medical Association guidelines. Hippocampi were isolated and sliced at 400 μm on a McIlwain Tissue Slicer (Brinkmann). Slices were placed in Hank’s balanced salt solution (HBSS) supplemented with glucose (27 mM, Sigma) before transfer to 0.4 μm Millicell membranes (Millipore) placed in 6well culture dishes. Growth media included 50% Minimal Essential Media, 25% normal horse serum, and 25% HBSS supplemented with 8 mM HEPES and 27 mM glucose. All cell culture media components were obtained from Life Technologies except where noted. Cultures were incubated at 35°C, 5% CO_2_ and 90% humidity for 5 days before returning to 36.5°C for the remainder of the 10-12 days before use.

### Oxygen-Glucose Deprivation (OGD)

HOTCs with normal architecture were exposed to anoxic and hypoglycemic conditions. Anoxic experiments were performed with Hanks Balanced Salt Solution (HBSS) which was pre-equilibrated to remove residual oxygen. HBSS was first bubbled with N_2_ outside of an anaerobic chamber (Coy Laboratories) for 30-60 minutes then transferred to plates which were placed in the chamber for another 4 hours. Slices were rinsed in and then placed in glucose-free HBSS and transferred to the pre-equilibrated media in the anaerobic chamber for 45 minutes at 36.5°C. Atmospheric conditions of the chamber included 85% N_2_, 10% CO_2_, and 5% H_2_. Anoxic conditions within the chamber were confirmed by an oxygen sensor (Coy Laboratories). After OGD treatment, HOTCs were returned to normal culture conditions for various times before harvest. Control slices were treated with HBSS containing glucose and left in normoxic conditions in a 36.5°C incubator.

### Cobalt Chloride Treatment

HOTCs were grown as above and subjected to CoCl_2_ treatment at 10-12 days after plating. Cobalt Chloride hexahydrate (CoCl_2_-6H_2_0, Sigma) was diluted in distilled water and diluted into HBSS supplemented media (as above) to the appropriate concentration (30-500 μM). Once HOTCs were placed in CoCl_2_, they were returned to a 36.5°C incubator until time of RNA harvest.

### RNA isolation

Total cellular RNA from HOTCs on multiple pooled plates was harvested from treated or control HOTCs using the RNeasy RNA Isolation Kit (Qiagen). RNA quality and quantity was analyzed with spectrophotometry and gel electrophoresis.

### Quantitative PCR

Quantitative-PCR experiments were performed using the StepOne Real-Time PCR System (Applied Biosystems). Briefly, isolated total RNA from slices was converted to double stranded cDNA using the High Capacity cDNA Reverse Transcription Kit (Life Technologies). Input cDNA was used at 2-50 ng per reaction as determined by studies to determine the linear ranges of amplification. Comparative C_t_ studies were performed using Taqman gene expression assays for Vegf, Epo, Hif1α, Epas1/Hif2α, Hif3α, Arnt/Hif1β, and Arnt2/Hif2β using the manufacturer’s cycling parameters. Normalization was performed using the endogenous control Hprt.

### Propidium Iodide

The toxicity of CoCl_2_ treatment on HOTCs was assayed using propidium iodide (PI), a fluorescent dye which can enter cells whose membranes are not intact and bind to the nuclear DNA [32, 33]. PI was added to CoCl_2_ containing media to a final concentration of 50 μM for 60 minutes before microscopy. An Olympus BX51WI upright microscope with Hammatsu ORCA-ER camera was used for taking bright field and fluorescent images. General slice health was assayed at the bright field level. Images of HOTCs were analyzed for PI intensity and compared to control slices as previously described [34].

### Nonquantitative RT-PCR

The splicing of Hif3α mRNA transcripts occurs in some systems when exposed to hypoxia [35–37]. Three sets of mouse Hif3α specific primers were used to analyze splicing of IPAS (Hif3α4 in humans) in our OGD and CoCl_2_ treated cultures [36]. Total RNA from slices was converted to double stranded cDNA using the High Capacity cDNA Reverse Transcription Kit (Life Technologies). An Applied Biosystems GeneAmp PCR System 9700 machine was used for amplification.

### Statistical Analysis

Statistical analyses were performed using GraphPad Prism software. Comparative ΔΔCt values were normalized to the reference gene HPRT (NRQ). Data shown are mean NRQ of 3 or more independent experiments. In experiments exhibiting decreases in transcription, the NRQ values were subjected to log (10) transformation [38] before one-way or two-way analysis of variance (ANOVA) tests were used to analyze the data, followed by post-hoc comparisons using the Dunnett’s Multiple Comparison Test. The level of significance was set at p<0.05 and results presented as mean +/− S.E.M.

## Results

### OGD shows time dependent increases in HIF target gene Vegf, but not Epo

HOTCs were subjected to anoxic conditions for 45 minutes in a glucose-free media (OGD) and assessed for induction of the HIF pathway by examining expression of two genes, Vegf and Epo, known to be regulated by hypoxia. QPCR on mRNA isolated from slices at 5 and 24 hours post insult showed increases in Vegf but not Epo mRNA levels as compared to control (p<0.01; Fig 1a). By 24 hours, Vegf expression returned to baseline (Fig 1a). While Epo gene expression trended towards an increase at 24 hours, it was not statistically significant (Fig 1b).

**Figure 1.**
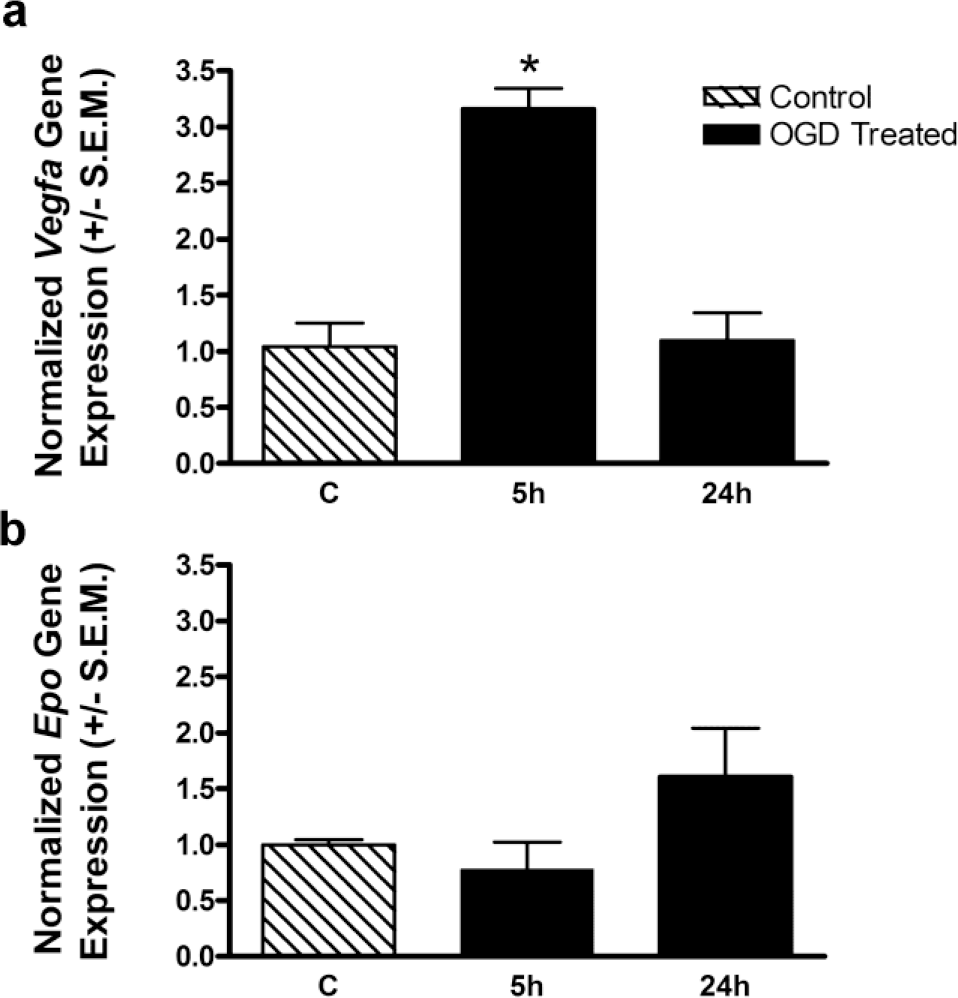
VEGF but not EPO transcripts are increased after OGD treatment. HOTCs were exposed to 45 min OGD, returned to normoxic conditions and RNA was harvested either 5 h or 24 h after insult. Time dependent induction of VEGF (a) and EPO mRNA expression (b) was assessed by q-PCR. Data are normalized to endogenous control Hprt and are shown as the mean of experiments performed in triplicate as fold induction above control. Control slices (c) remained in normoxic conditions for the duration of the experiment.

**Fig 1. Time dependent induction of Vegf (a) and Epo (b) in slices exposed to OGD**. Slices were exposed to 45 min OGD and RNA was harvested either 5 h or 24h after insult. Data are normalized to endogenous control Hprt. N =3-5 independent experiments, * p < .01 compared to untreated (control) slices.

### HOTCs respond to CoCl_2_, a hypoxic mimetic, in a time and concentration dependent manner

To examine the effects of HIF pathway activation on gene activation using a chronic model, HOTCs were treated with CoCl_2_ for comparison with OGD. CoCl_2_ allows HIF formation by inhibiting PHD and VHL activity [6, 7]. CoCl_2_ treatment of HOTCs to induce the HIF pathway has not been published; therefore, we used treatment concentrations used for dissociated cultures to determine starting concentrations [39, 40]. Preliminary experiments amplified both Vegf and Epo targets. However, Vegf expression was more robust and has been shown to be expressed very rapidly after HIF activation and so was used to determine the time course and concentration of future experiments [41].

Figure 2 shows the time and concentration dependent induction of Vegf mRNA after CoCl_2_ treatment. Two factor analysis of variance indicated a significant overall effect of concentration. Lower doses of CoCl_2_ (30 and 100 μM) showed more modest increase of Vegf expression that is both time and concentration dependent. At higher concentrations of CoCl_2_, only a 5 hour exposure time was needed to significantly increase the expression of Vegf mRNA as compared to control (Fig 2). Expression remained elevated at 24h with no significant change from 5 hours. Thus, the expression of Vegf mRNA showed both dose and time dependent expression that plateaus at higher concentrations after treatment with CoCl_2_.

**Figure 2.**
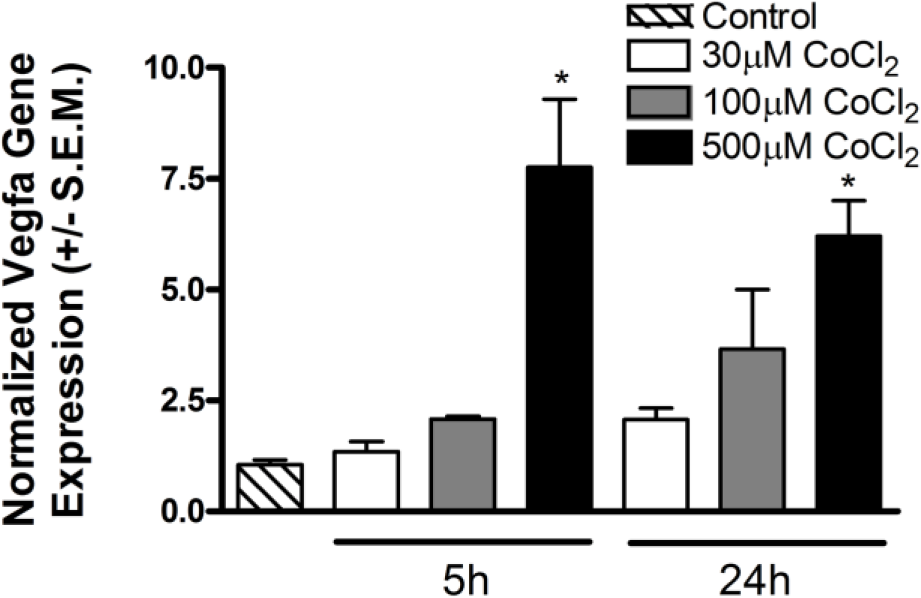
VEGF transcripts are increased after treatment with high doses of CoCl_2_. HOTCs were treated with indicated concentration of CoCl_2_ (30-500 mM) and RNA harvested either 5 or 24 h later. Q-PCR analysis of VEGF mRNA expression was performed for comparison with control slices. Data are normalized to endogenous control Hprt and are shown as the mean of experiments performed in triplicate.

**Fig 2. Time and concentration dependent induction of Vegf after treatment with cobalt chloride**. Slices were treated with indicated concentration of CoCl_2_ and RNA was harvested either 5 or 24h later. N = 3-7 independent experiments, *p < 0.001 compared to control.

### CoCl_2_ toxicity in HOTCs is time and concentration dependent

CoCl_2_ toxicity on HOTCs was assayed using microscopy to examine different aspects of cell integrity. Wells of 4-6 HOTC slices were exposed to various concentrations of CoCl_2_ (30-500μM) for 5 and 24 hours for comparison with control treated HOTCs. Light microscopy was used to examine changes associated with the health of the slice, including color and morphology. In addition, propidium iodide (PI) staining was performed to examine the presence of dying cells. At 30 and 100 μM CoCl_2_ (5 and 24 hours), HOTCs were clear, with easily distinguished morphological layers and little PI staining (data not shown). Similar results were found with 500 μM treated slices at 5 hours after treatment. In contrast, by 24 hours, 500 μM CoCl_2_ treated slices appeared brown, condensed and lacked normal architecture. PI staining of 500 μM treated HOTCS showed a diffuse, non-layer specific increase in dying cells that was statistically significant at 24h compared to control (p<0.05, Fig 3b). In contrast, at 24-hour post-OGD, HOTCs show a CA1 specific increase in PI ([32] and unpublished data).

**Figure 3.**
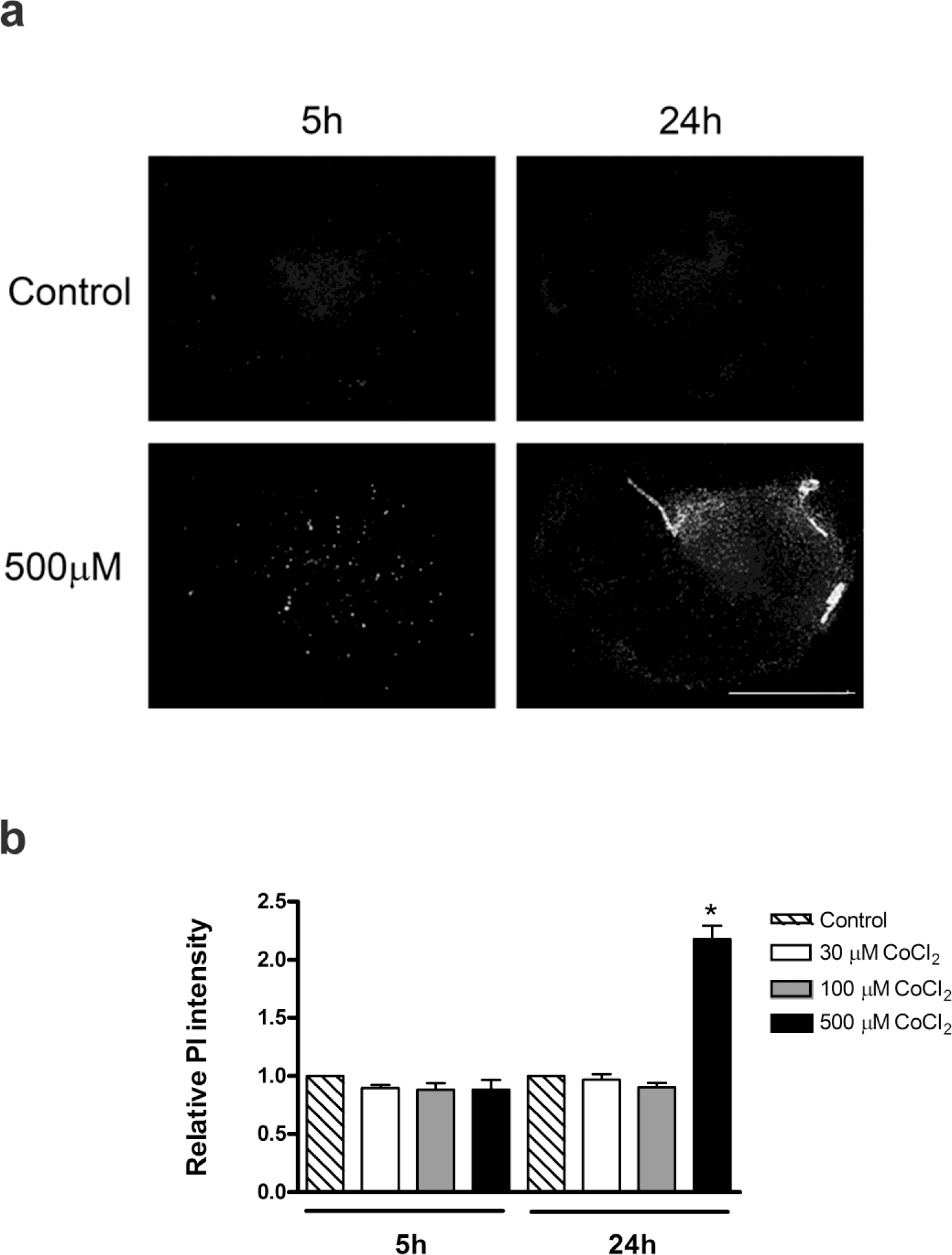
CoCl_2_ causes degenerative changes to HOTCs at high concentrations after chronic treatment. Slices were exposed to indicated concentration of CoCl_2_ (30-500uM) for 5 or 24 hours. HOTCs were stained with propidium iodide for 60 minutes before assaying for penetrance of the dye. Increased staining indicative of cell death can be seen in individual slices treated with CoCl_2_, but not control slices (a). Quantitation of staining intensity revealed a significant increase in cell death after treatment with 500 mM cobalt chloride (b).

**Fig 3. Propidium Iodide staining in slices treated with cobalt chloride**. Slices were exposed to indicated concentration of CoCl_2_, followed by incubation with PI after 5 or 24 hr. Increased staining, indicative of cell death, can be seen in individual slices (a). Quantitation of staining intensity revealed a significant increase in cell death 24h after treatment with 500 μM CoCl_2_ (* p < .05, n = 5-10).

### Differential temporal expression of Vegf and Epo mRNA occurs with 30 and 500 μM CoCl_2_ time course treatments

In Figure 2, we saw a time and concentration dependent expression of Vegf mRNA with HIF activation by CoCl_2_. To further investigate the dose-specific temporal changes in expression of HIF targets after CoCl_2_ treatment, Vegf and Epo gene expression was assayed at additional time points using a high and low dose of CoCl_2_ (30 and 500 μM). Treated slices were collected in two hour increments between 5 and 24 hours to ensure that transient changes in expression were not missed. This time frame was chosen because detectable increases in Vegf mRNA expression denoting induction of the HIF pathway do not begin until at least 5 hours after treatment using 500 μM CoCl_2_ (unpublished results). Further, significant neuronal death occurs in HOTCs after OGD and CoCl_2_ by 24 hours (Fig 3) [34]. Vefg and Epo mRNA expression was assayed at 11 time points (5, 7, 9, 11, 13, 15, 17, 19, 21, and 24 hours after CoCl_2_ treatment) for comparison to control.

Vegf expression increased significantly as compared to control in 30μM CoCl_2_ treated slices after 15 and 17 hours of treatment, but at no other time points (p<0.05; Fig 4a). Unlike OGD treatment (Fig 1), 30 μM CoCl_2_ treatment did not provide an early increase in Vegf mRNA expression (Fig 4a, open bars). However, using 500μM CoCl_2_, significant increases in Vegf expression were seen at every time point from 5 to 24 hours (p<0.05; Fig 4a, solid bars). Epo mRNA expression differed quite significantly. While Vegf mRNA induction was greater at 500 μM than at 30 μM, Epo mRNA expression showed the inverse trend. Epo expression with 30 μM CoCl_2_ increased during the first 15 hours to a significant level at 17-24 hours (Fig 4b, p<0.05). In contrast, there was an early but transient equivalent expression of Epo at 5 hours with 500 μM CoCl_2_ that declined to insignificant levels for the duration of the treatment (Fig 4b, solid bars).

**Figure 4.**
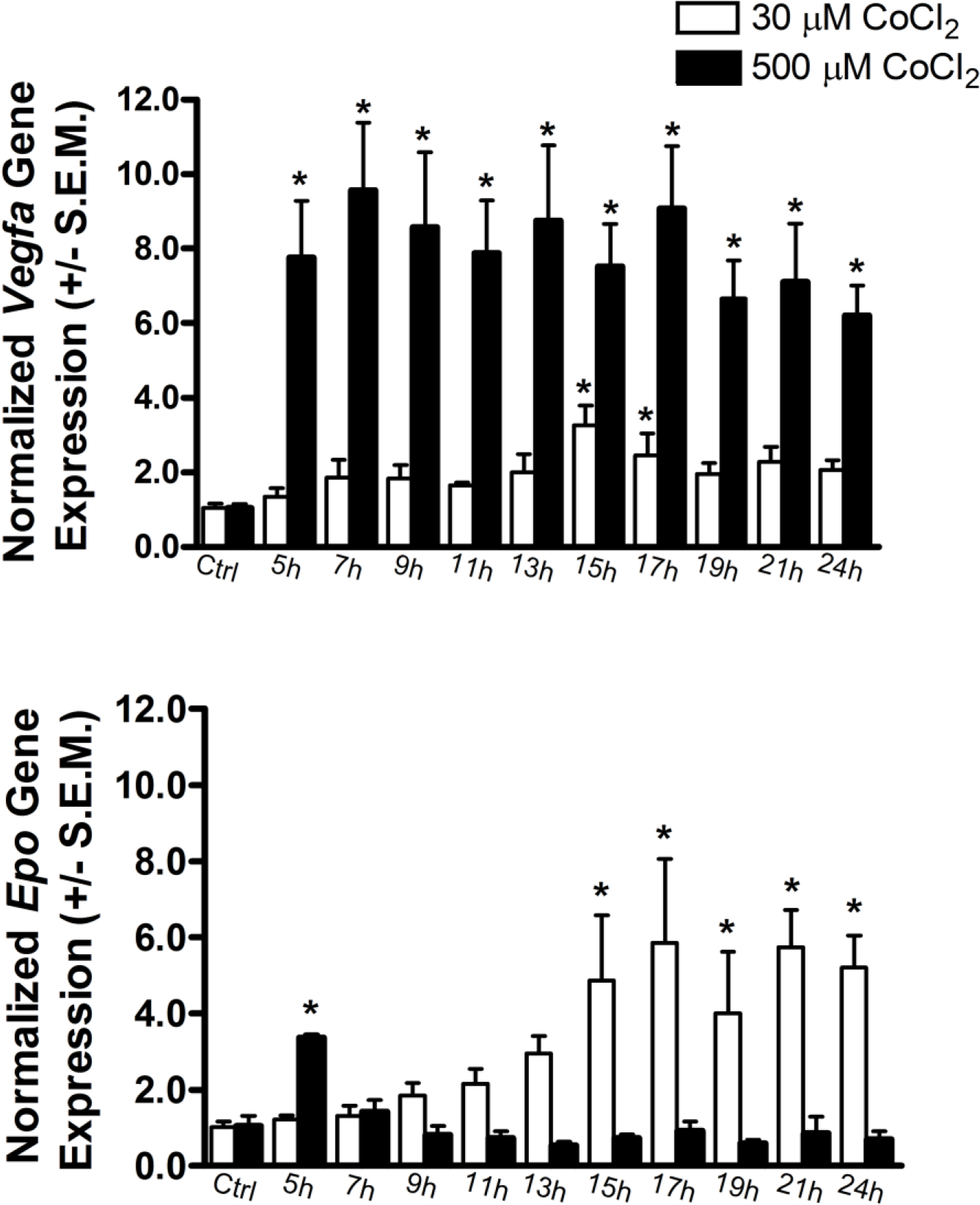
High and low concentrations of CoCl_2_ regulate VEGF and EPO mRNA gene expression differently. HOTCs were treated with indicated concentration of CoCl_2_ (30 or 500 mM) and RNA harvested at 2 hour intervals between 5 and 24 hours. Q-PCR analysis of VEGF and EPO mRNA expression was performed for comparison with control slices. Data are normalized to endogenous control Hprt and are shown as the mean of experiments performed in triplicate. Shown is the time course of changes in expression of VEGF (a) and EPO (b) following treatment with low or high concentrations of CoCl_2_. Expression is represented as fold induction above untreated control.

**Fig 4. Time course of changes in expression of Vegf (top) and Epo (bottom) following treatment with low or high concentrations of CoCl_2_**. Values are normalized to Hprt and expressed as fold induction above control (untreated). Each time point represents 3-6 independent experiments, * indicates significant difference p < 0.05 from control.

### Hifα subunits are selectively affected by induction of OGD

QPCR was performed to assess mRNA expression changes that occur in the structural subunits of HIF after induction of the HIF pathway. Three genes encode structural alpha subunits that encode proteins involved in HIF complex formation (Hif1α, Epas1/Hif2α, Hif3α). Slices were exposed to OGD for 45 minutes and harvested at 5 and 24 hours for comparison to control slices. An additional 3 hour time point was also examined for the Hif1α target to elucidate trends that we saw occurring at 5 hours (data not shown). QPCR shows that Hif1α expression increased at 5 hours (p<0.05) and returned to baseline by 24 hours (Fig 5a). Hif2α and Hif3α expression were also examined. Fig 5b shows expression of Hif2α, which was not significantly different at any time examined. Hif3α expression showed a trend towards decreased expression at 5 hours post treatment that was not significant, but similar to control levels by 24 hours (Fig 5c).

**Figure 5.**
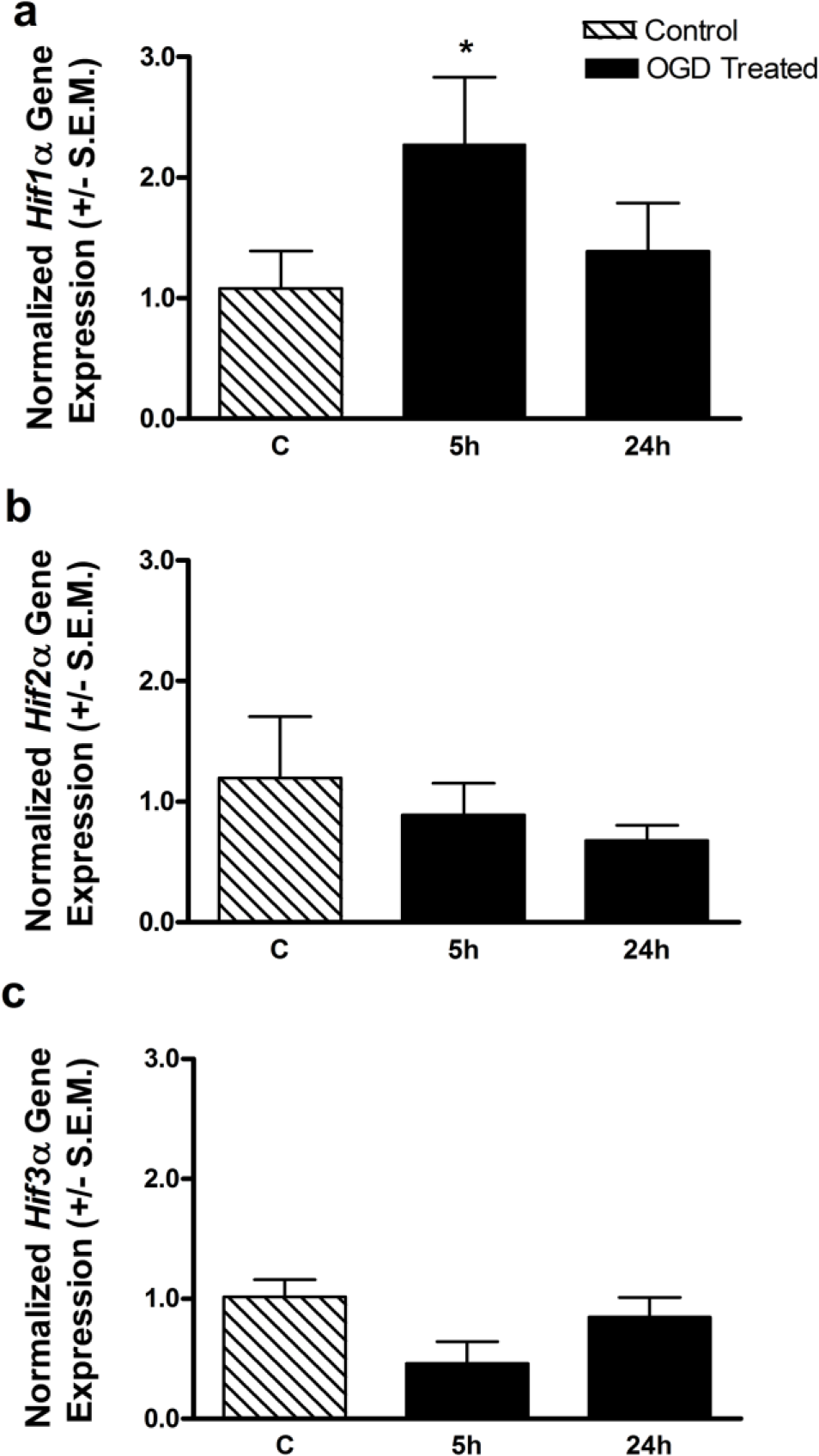
OGD differentially regulates mRNA gene expression of HIF alpha subunits. HOTCs were treated with OGD and RNA harvested at 5 and 24 hours after insult. Q-PCR analysis of HIF1A (a), HIF2A (b) and HIF3A (c) mRNA expression was performed for comparison with control slices. Data are normalized to endogenous control Hprt and are shown as the mean of experiments performed in triplicate. Expression is represented as fold induction above untreated control.

**Fig 5. Expression of Hif subunits (Hif1α, Hif2α, Hif3α) in slices exposed to 45 min OGD**. Expression data is normalized to endogenous control Hprt. *p < 0.05 compared to control (C).

### The expression of HIFα subunits is selectively affected by CoCl_2_

Slices were exposed to two concentrations of CoCl_2_ (30 and 500 μM) at 11 time points, starting at 5 hours and ending at 24 hours, and assayed for changes in Hifα expression as compared to control. Hif1α expression decreased with 500 μM CoCl_2_ at later time points (19-21 hours), while 30 μM CoCl_2_ did not induce significant changes from control (p<0.05, Fig 6a). Hif2α and Hif3α expression was also examined using the same time course (Fig 6b, c). Hif2α expression did not change at either concentration of CoCl_2_ (Fig 6b). Hif3α expression was significantly decreased by treatment with 500 μM CoCl_2_ at all times examined (p<0.05), while not changed with 30 μM treatment (Fig 6c).

**Figure 6.**
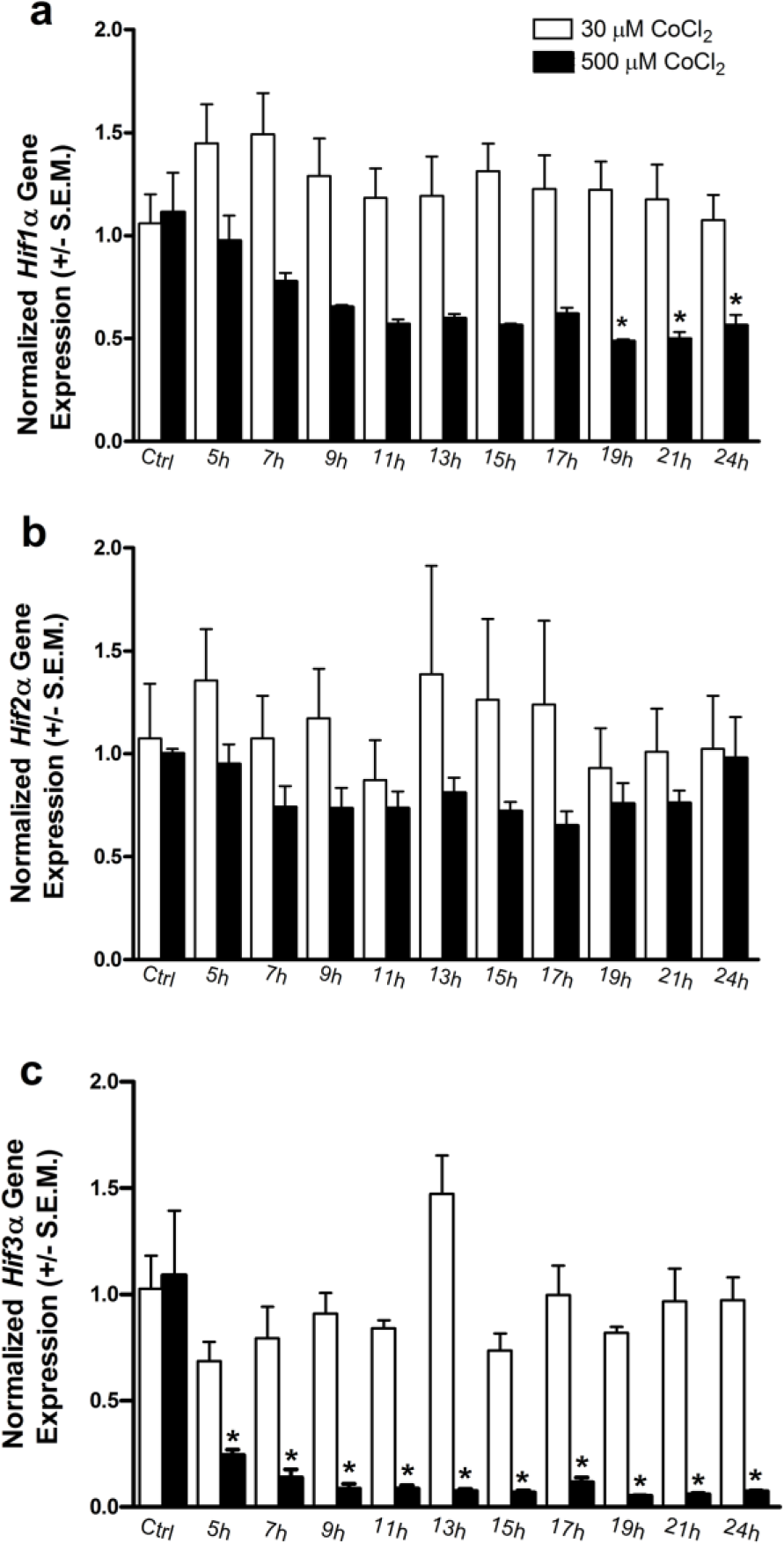
CoCl_2_ decreases mRNA gene expression of HIF3A subunits. HOTCs were treated with indicated concentration of CoCl_2_ (30 or 500 mM) and RNA harvested at 2 hour intervals between 5 and 24 hours. Q-PCR analysis of HIF1A (a), HIF2A (b) and HIF3A (c) mRNA expression was performed for comparison with control slices. Data are normalized to endogenous control Hprt and are shown as the mean of experiments performed in triplicate. Expression is represented as fold induction above untreated control.

**Fig 6. Time course of expression of Hif subunits in slices treated with low or high concentrations of CoCl_2_**. Expression is normalized to endogenous control Hprt and compared to control slices. Each result represents 3-9 independent experiments. * p < 0.05

### Hif3α splice variants are not found after OGD or CoCl_2_treatments in HOTCs

Hif3α is a dynamic protein that can be alternatively spliced after hypoxic conditions. Splice variant expression has been reported after hypoxia in cerebellum, cornea, heart, lung and kidney [35, 36, 42]. In our system Hif3α splice variants were not detectable by the Taqman gene assay selected for QPCR. Thus, non-quantitative PCR was used to try to detect splice variant expression. HIF3α primers designed to non-variable regions of the gene were used as positive controls. Amplification of HIF3α was detected in the control and treated cultures (both OGD and CoC1_2_ at 30 and 500 μM). However, Hif3α splice variant specific primers did not detect expression in control or treated slices, thus we have no evidence of their expression in our culture system [36].

### HIFβ subunits are not changed by OGD treatment

QPCR was performed to assess the mRNA expression changes that occur in the beta structural subunits of HIF after CoCl_2_ or OGD treatments. Two genes encode structural beta subunits that encode proteins involved in HIF complex formation (Hif1β/ARNT, Hif2β/ARNT). OGD treated slices did not show significant Hif1β or Hif2β changes when assayed at 5 and 24 hours after treatment (Fig 7a, b). However, CoCl_2_ treated slices did show decreases in Hif1β and Hif2β expression when assayed by QPCR from 7-24 hours (Fig 7c, d).

**Figure 7.**
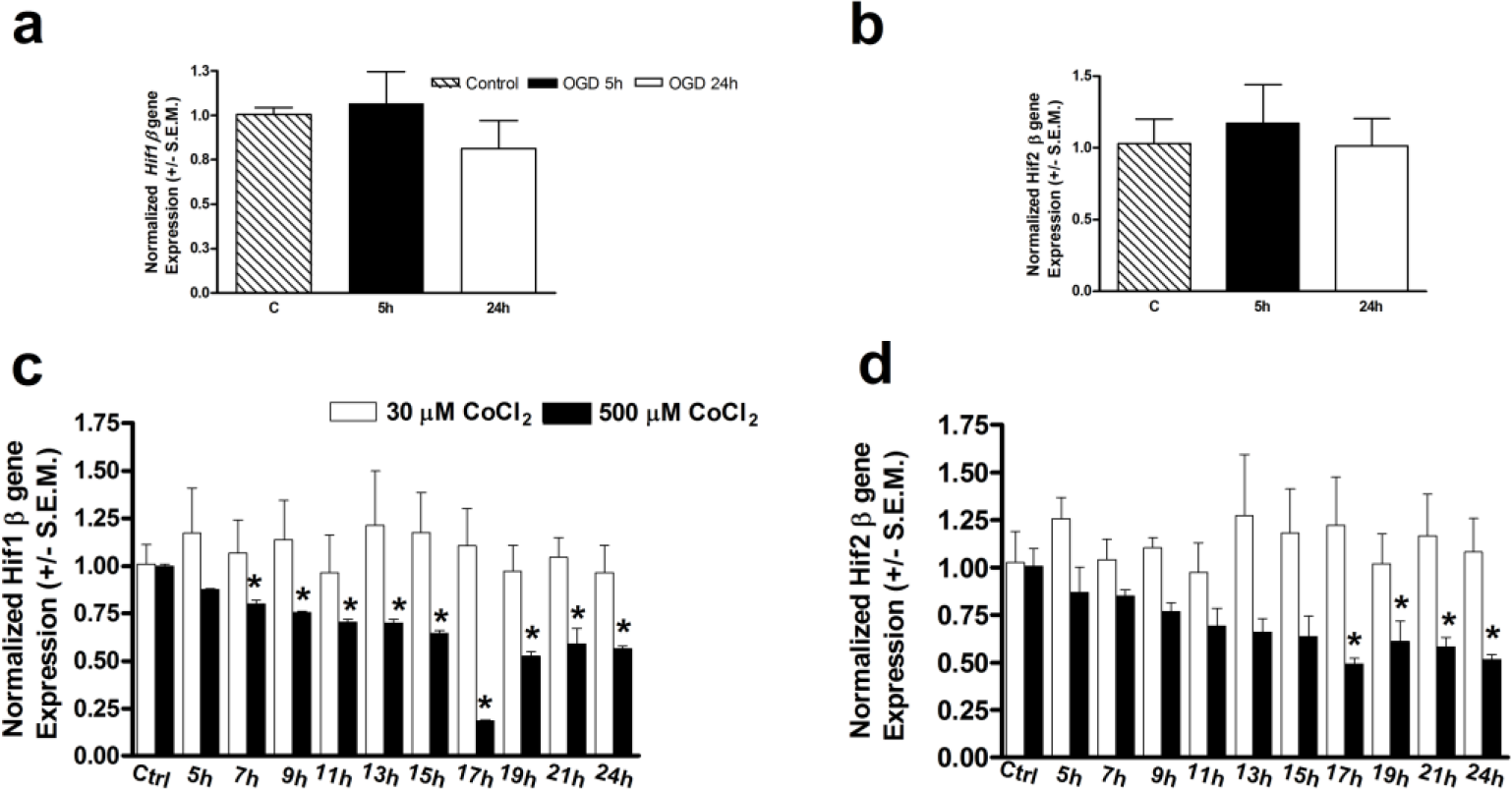
HIF beta subunit mRNA expression is unaltered by both OGD and CoCl_2_ treatments. HOTCs were treated with OGD and RNA harvested at 5 and 24 hours after insult (a,b). HOTCs were treated with indicated concentration of CoCl_2_ (30 or 500 mM) and RNA harvested at 2 hour intervals between 5 and 24 hours (c,d). Q-PCR analysis of HIF1B (a, c), HIF2B (b, d) mRNA expression was performed for comparison with control slices. Data are normalized to endogenous control Hprt and are shown as the mean of experiments performed in triplicate. Expression is represented as fold induction above untreated control.

**Fig 7. Hifβ subunits are not affected by OGD**. Slices were exposed to 45 min OGD (a, b) and expression of Hif1β (a) and Hif2β (b) following 5 or 24h was determined. Values are fold induction compared to control cultures. In separate experiments (c and d), slices were exposed to 30 μM (open bars) or 500 μM CoCl_2_ (black bars) and expression of Hif1β (c) and Hif2β (d) was measured at time points from 0-24h. * p < 0.05.

## Discussion

One of the foremost challenges in clinical neurology is the treatment of ischemia in the nervous system, such as that found in stroke and traumatic injuries to the brain and spinal cord. To develop therapies that can provide clinical treatment for patients experiencing reduced oxygen levels affecting the nervous system, one must first understand the molecular mechanisms involved in these processes. For the purposes of this manuscript, we have focused on changes occurring during the first 24 hours after insult in our model systems. Ischemic conditions are caused by the reduction of both oxygen and glucose. Oxygen deprivation studies have been done *in vivo* and *in vitro* to examine the effects of ischemia and hypoxia in acute and chronic settings [19, 21, 27, 28, 34, 43]. In each system, the common factor is that a reduction in oxygen causes activation of the HIF pathway, a heterodimeric system containing a labile HIFα and stable HIFβ subunit. The magnitude of HIF activation is dependent on the extent of oxygen deprivation and is maximized at O_2_ levels of 0.5%, a number similar to that seen during ischemia [43]. Glucose may also play a role in the degree of activation of the HIF pathway. In neuroblastoma cells, the presence of glucose gave greater Hif1α protein stabilization as well as increased transcription of HIF target genes [44]. They concluded that the absence of glucose, such as found in oxygen glucose deprivation (OGD), dampens HIFα stabilization and subsequent target gene expression [44]. However, they did not test the expression of specific genes. Thus, the role of glucose in HIF target gene induction may yet be defined better here.

In our studies, the HIF pathway was activated in two ways, with either a transient oxygen decrease accompanied by a loss of glucose (OGD), or by pharmacologically mimicking hypoxia with CoCl_2_. While OGD is a commonly used method in HOTCs, the use of CoCl_2_ with HOTCs has not been reported in the literature before now. CoCl_2_ has been used in dissociated cultures to induce hypoxic conditions [45]. Further, *in vitro* studies have reported CoCl_2_ dependent increases in the expression of HIF dependent genes [18, 24, 40]. However, these models are different in their duration, degree of oxygen deprivation and presence of glucose. While looking at similar start and end points, the CoCl_2_ model mimics chronic hypoxia with glucose while OGD mimics acute hypoxia with reperfusion. Thus, these two models have the potential to give us different information about HIF induction in the HOTC and the role of glucose.

### HIF pathway induction of target genes

To characterize the induction of target genes in the HIF pathway in HOTCs, Vegf and Epo expression was examined. Both Vegf and Epo mRNA expression is mediated by the HIF pathway and can be induced by hypoxic conditions [3–5, 46–48]. These molecules are transcribed after hypoxic exposure and allow a physiological adaptation to a loss of oxygen through compensatory mechanisms [49].

In our systems, Vegf mRNA expression was increased by both OGD and CoCl_2_, with a dose and time dependence. Only chronic application of 500 μM CoCl_2_ with glucose increased expression significantly at all time points examined. Further, OGD induced expression was increased only 3.1 fold versus 7.5 fold following 5 hours of treatment with 500 μM CoCl_2._ Thus, the absence of glucose in the OGD condition may be dampening this response as suggested previously [44] Vegf mRNA expression also shows a clear CoCl_2_ dose dependence in dissociated hippocampal cultures [50]. While oxygen reduction is a large player in the increase of Vegf mRNA, additional cellular factors that could induce Vegf expression have also been described but are not well understood in the HOTC, [5, 51, 52]. Changes in expression of several Vegf inducers (platelet derived growth factor [PDGF], signal transducer and activator of transcription [STAT] and cyclooxygenase −2 [Cox-2]) have been reported to be altered in separate but non-overlapping ischemic microarray panels [5, 52, 53]. Presently, there is no evidence that of any of these potential Vegf inducers are active in our system and thus the significance in the HOTC is yet unknown. It is also unknown whether there is a change in the stability of the Vegf transcripts due to reduced oxygen conditions as seen in some hypoxic models [4, 49, 54]. Thus, there may be additional regulatory pathways that may play into the expression patterns that we are seeing.

Epo gene expression was also examined using OGD and CoCl_2_ treatments. In our HOTC model, OGD treatment did not produce significant increases in Epo mRNA expression at any time point while CoCl_2_ treatment did. Previous *in vivo* studies in cerebellum and cerebrum have shown Epo mRNA levels to be significantly elevated after hypoxia [55–57]. Epo mRNA expression has not been reported following oxygen and glucose deprivation in HOTC. These differences may again be linked to the total absence of glucose in the OGD condition. CoCl_2_ induced expression of Epo was again dose dependent, but in inverse relation to Vegf. Low levels of CoCl_2_ induced a delayed increase in Epo (30 μM) at 15-24 hours, while 500 μM CoCl_2_ rapidly but transiently induced Epo at 5 hours. This differs from our results in cultured astrocytes, where low concentrations of cobalt induced transcription of EPO after 5 hours of treatment [50].

There may be several factors other than the absence of glucose which contribute to transcriptional regulation differences. During chronic hypoxia, Epo transcripts are down regulated due to negative feedback in the HIF system created by increased transcription of the PHD-2 and PHD-3 genes [46]. In addition, several studies have reported that EPO induction is Hif2α dependent [58, 59]. In cancer cell cultures in which hypoxia induced EPO expression is Hif2α dependent, CoCl_2_ was unable to induce Epo mRNA expression [58]. Thus, unlike hypoxia, it may be that CoCl_2_ activates HIF2α dependent expression of Epo in a delayed and sustained fashion at low concentrations and that higher CoCl_2_ accelerates the inhibition of HIF2α protein through feedback mechanisms. This may explain why lower concentrations of CoCl_2_ (30 μM), produce greater expression of Epo than 500 μM CoCl_2_. Further, while Epo induction has been primarily described as HIF dependent, Gata-2 and NF-Kβ have been shown to decrease Epo expression by binding to the promoter region of the Epo gene [60]. In sum, Epo has a very complex and tightly regulated expression at the RNA level whose altered expression is evident in many types of systems.

### HIF structural subunits are differentially regulated by OGD and CoCl_2_

To further understand the self-regulation of the HIF pathway through feedback mechanisms (direct or indirect), we also assessed the mRNA expression of the alpha and beta subunits of HIF. Previous literature has reported that beta subunits are hypoxia resistant with little regulation, which was confirmed in our OGD but not CoCl_2_ experiments [19, 27, 28]. In fact, we observed decreased expression of Hif1β and Hif2β with chronic CoCl_2_ treatment.

In contrast, HIF alpha subunits were not only regulated by OGD and CoCl_2_, but differentially by subunit. In the literature, there are conflicting reports about Hif1α mRNA expression. Transcriptional upregulation of Hif1α in the brain has been shown *in vivo* after sustained hypoxic treatment or ischemic treatment in some systems [19, 21, 23, 28]. However, conflicting *in vivo* studies using hypoxia alone have not reported changes in Hif1α gene expression [19, 27]. In our OGD system, Hif1α mRNA levels were increased at 5 hours, which declined to baseline by 24 hours. Using similar HOTC methods, Lushnikova’s group found that individual hippocampal neurons (in CA1 and CA3) showed different responses to OGD 4 hours after OGD end[21]. However, neither cell type showed an increase in Hif1α transcripts at 4 hours post-OGD. Our data indicates a significant Hif1α transcript increase at 5 hours post-OGD, however this mRNA is pooled from the entire slice and thus there may be other cell types in the HOTC in which Hif1α is being elevated. Based on our results and those in the literature, a loss of glucose, such as that seen with OGD, may promote the cellular pathways which are responsible for induction of Hif1α gene expression. Hif1α induction has also been linked to upstream regulatory pathways involving NF-kB, which is induced during hypoxia [23, 61]. From Lushnikova’s experiments, it is clear that different cell types may have different expression of Hif1α which may ultimately affect their ability to survive hypoxic or ischemic insults. CA1 neurons are very sensitive to ischemic insult, and this may due in part to their inability to transcribe Hif1α, a critical component of the HIF protein complex.

In contrast to OGD, treatment of HOTCs with CoCl_2_ (500 μM) produced modest down regulation of Hif1α mRNA at much later times. A transient decrease of Hif1α was seen at 19 and 21 hours CoCl_2_ treatment. CoCl_2_ regulation of Hif1 mRNAs has not been reported in the literature until now. Due to the differences in the time windows examined, it is difficult to know if OGD also down regulates Hif1α at 19 and 21 hours similarly to CoCl_2_. The down regulation of both Hifβ and Hifα subunits with chronic CoCl_2_ treatment may be indicative of compensatory down-regulation in the Hif pathway in the event of chronic insult. It is evident that the regulation is a multi-factorial process, and looks very different in acute versus chronic oxygen depletion, with further differences being made with the absence of glucose.

### OGD down regulates Hif3α transcripts

The Hif3α gene is the only Hif structural subunit that contains a hypoxia response element (HRE) and thus has a potential direct regulation by HIF-1 [37]. Hif3α expression is variable, depending on the system being studied. Within the hippocampus, Hif3α expression levels are decreased in CA1 neurons after OGD [21]. This is consistent with our OGD results from the whole slice, which showed a nonsignificant decrease in Hif3α mRNA at 5 hours. Similarly, CoCl_2_ treatment with 500 μM showed a drastic reduction in Hif3α that was sustained at all time points analyzed.

The existence of alternatively spliced variants of the HIF3α gene, encoding several isoforms of HIF3α, has been reported and further complicates our understanding of its function [35, 36, 62]. A loss of oxygen causes the selective alternative splicing of the inhibitory PAS (Per/Arnt/Sim; IPAS) variant, with a concomitant reduction in non-spliced Hif3α gene expression [35, 36, 62]. IPAS has an HRE in its upstream region which is targeted by HIF1α and its protein serves as a dominant negative feedback regulator which inhibits HIF gene transactivation [37]. Normoxic IPAS expression has been described in cerebellum and corneal epithelium; however IPAS splice variant generation has not been described in other types of neurons during hypoxic conditions [35]. Using RT-PCR, we were unable to detect expression of any of the splice variants, including IPAS, with either OGD or CoCl_2_ treatments (data not shown). The lack of IPAS expression is supported by the observation that IPAS and Vegf expression are inversely related in other systems [35]. Thus, the IPAS splice variant may not be induced in hippocampal cells in our culture system or perhaps in neurons in general. However, it remains to be seen why Hif3α mRNA transcripts are downregulated in our system if there is not a concomitant increase in IPAS transcription. It may be that the Hif3α protein can act as a transcriptional regulator of HIF1α and HIF2α in this system (similarly to IPAS) which is feeding back to decrease its own transcription. Additional studies will be helpful in understanding the expression of IPAS in neurons of the brain.

## Conclusions

Transcriptional regulation is a hallmark of exposure to a reduction in oxygen. Here, results show a side by side comparison of the induction of expression of molecules involved in or targets of the HIF pathway using OGD and CoCl_2_. While Vegf expression was induced by both treatments, differential increases in transcription of Epo (only with CoCl_2_) and Hif1α (only with OGD) were seen. Thus, these models have the ability to elucidate the transcriptional activation of different genes in the same HOTC model. OGD and CoCl_2_ also differentially regulate HIF structural subunits, which plays into the overall functioning of the HIF system during periods of altered oxygen. Further, while the scope of this work focused on changes occurring during the first 24 hours after insult, future studies would benefit from extending this time frame. Additional studies will need to elucidate the potential role of glucose in CoCl_2_ induction of transcription and the importance of Hif3α regulation in the hippocampus.

## Acknowledgements

The authors would like to thank Jeong-Woo Seo for laboratory support and Jessica Snell for animal care.

